# Temple-measured postprandial cardiovascular responses under free-living conditions

**DOI:** 10.64898/2026.11.23.768469

**Authors:** Deland Liu, Michael P. Snyder, Sachin Nadig

## Abstract

Eating perturbs cardiovascular homeostasis: splanchnic vasodilation after a meal is offset by autonomic adjustments that raise heart rate, and both the magnitude and duration of this response can depend on meal size and composition. Measuring these responses in daily life requires knowing when each meal begins and ends, which self-report captures poorly. The temple region allows both to be measured with a single device: chewing-related motion over the temporalis can mark meal onset and offset, and photoplethysmography (PPG) at the superficial temporal artery (STA) records the cardiovascular response. The STA also supplies a vascular bed whose postprandial behavior has not been well characterized. Using a temple-worn wearable under free-living conditions, we asked (1) whether established postprandial heart rate (HR) responses can be observed from automatically detected meals, and (2) how PPG pulse morphology at the STA changes after a meal. One meal from each of 100 participants in free-living conditions was analyzed, comparing a 15-min pre-meal baseline with the 120 min after eating, with nutrient content estimated from meal photographs. HR rose approximately 2.6 bpm on average in the first 15 min after eating and approached baseline within two hours. The postprandial rise in HR increased with caloric content. HR remained elevated two hours after the most sodium-dense meals, whereas higher fiber density was associated with a larger rise only during the first 15 minutes. The heart-rate increase per 100 kcal was greater in participants with higher BMI. In addition, the PPG morphology index *A_d_/A_a_*, a second-derivative PPG marker of vascular tone and wave reflection, increased within 15 minutes of the meal and remained shifted at two hours at the temple region. This change persisted after adjustment for heart rate and is consistent with reduced wave reflection, possibly associated with vasodilation in vessels downstream of the aorta. These results support the feasibility of using a single temple-worn device to detect meals and measure the cardiac and vascular responses that follow in daily life, and identify the temple as a site for studying postprandial vascular changes.

## 1 Introduction

Eating imposes a recurring challenge on cardiovascular homeostasis: a meal typically perturbs the distribution of blood flow, and the circulation continuously adjusts to keep arterial pressure and organ perfusion stable [22]. The transition from the fasted to the postprandial state is marked by vasodilation of the gastrointestinal vasculature and a resulting increase in blood flow to support digestion and nutrient absorption [16, 30]. Because this vasodilation lowers systemic vascular resistance and shifts blood volume into the abdominal circulation, it could reduce arterial pressure if left uncompensated. Instead, in healthy individuals, blood pressure is generally maintained through an increase in cardiac output and vasoconstriction in non-splanchnic vascular beds [39, 37]. When these compensatory mechanisms fail, as in many older adults and patients with autonomic dysfunction, postprandial hypotension can result [19]. These hemodynamic adjustments are coordinated largely by the autonomic nervous system, through increased sympathetic outflow accompanied by reduced cardiac vagal tone [44, 13]. This autonomic shift is generally reflected in a rise in heart rate (HR) and a change in heart rate variability (HRV) [27, 42].

The magnitude and time course of postprandial hemodynamic changes can depend on both the energy content [37, 45] and the composition of a meal [39, 30]. Eating produces rise and recovery trend in heart rate (HR). In healthy adults, cardiac output increases after a meal through increases in both HR and stroke volume [23, 45], while total peripheral resistance falls [45]. HR has been shown to be higher around 30 min after a meal than both before eating and 110 min after eating [18], and after a small meal cardiac output returns close to baseline within 120 min [45]. The magnitude and duration of this response can scale with the calorie content of the meal. The maximum postprandial rise in HR was greater after 2 and 3 MJ meals than after a 1 MJ meal [37]. Across meals of 280, 560 and 840 kcal, HR increases were smaller after the 280 kcal meal at all time points, and smaller after the 560 kcal than the 840 kcal meal at 15, 30 and 120 min [1]. Studies that deliver glucose directly into the duodenum suggest the same load dependence: HR rose during infusions of 2 and 3 kcal/min but not 1 kcal/min [43]. Larger meals also prolong the response: cardiac output remained elevated 120 min after a large meal but had returned close to baseline after a small one [45], and superior mesenteric artery flow remained higher after larger meals for 90–120 min [37, 1]. Fewer studies have isolated individual components such as sodium and fiber, and studied their associations with postprandial HR response. Compared with a low-salt meal, a high-salt meal impairs brachial flow-mediated dilation [9] and increases the augmentation index [8], and eating salted snacks has been followed by increases in HR and cardiac output [25]. Viscous fiber slows gastric emptying and glucose absorption and attenuates the postprandial fall in blood pressure [20, 36], and adding guar to an intraduodenal glucose infusion attenuated the accompanying rise in HR [32]. Because sodium and fiber influence how blood vessels and blood pressure respond to a meal, they may also be associated with the postprandial HR response, although this has not been tested extensively in mixed meals. Beyond the cardiac response, eating also alters the arterial pulse itself. Eating has been associated with reduction in arterial wave reflection, the portion of each pressure pulse that is reflected from the peripheral arteries back toward the heart. This can be assessed by the augmentation index, which reflects how much the returning wave adds to central blood pressure, and it decreases after a meal [2], and after accounting for the accompanying changes in HR [41].

Monitoring of postprandial responses in free-living can be important for both clinical and healthy populations. For example, eating increases cardiac demand, which may be clinically relevant in people with coronary artery disease, whose capacity to meet that demand is limited. Postprandial angina occurs in up to 10% of patients with chronic ischemic heart disease and has been attributed to increased myocardial oxygen demand after eating, together with a redistribution of coronary blood flow away from regions supplied by stenosed arteries [7, 3, 5]. This demand can also be prolonged: in patients with coronary artery disease, cardiac output remained elevated two hours after a meal [24]. Heavy meals have further been associated with an increased risk of acute coronary syndrome [26]. For these patients, both the magnitude and the duration of the postprandial cardiac load therefore matter. Because HR might be an indicator of myocardial oxygen demand and can be measured continuously, tracking the magnitude and duration of postprandial HR elevation may help flag meals that impose an unusually high cardiac load. Conversely, a blunted HR response may signal impaired cardiovascular regulation, particularly in older adults and people with autonomic dysfunction (e.g., due to diabetes) [28]. The postprandial rise in HR is part of the compensatory response that maintains blood pressure as blood is redirected to the splanchnic circulation. When this compensation is impaired, blood pressure can fall, resulting in postprandial hypotension [19]. Detecting a consistently blunted HR response could therefore potentially prompt further clinical assessment. In healthy individuals, tracking postprandial HR responses over time may support healthier habits. Because larger meals tend to produce larger and more prolonged HR responses [45, 37, 1], such tracking could provide a rough, passive indication of meal size without manual food logging, which could suffer from poor long-term adherence [6]. It could also help users identify meals associated with unusually large or prolonged responses and adjust their routines accordingly. For example, it could reveal when a late meal leaves HR still elevated at bedtime, prompting users to eat dinner earlier or reduce its size. Similarly, because HR can remain elevated for some time after eating, tracking its return to baseline after different meals could help users judge how long to wait before strenuous exercise, potentially reducing gastrointestinal discomfort.

Monitoring postprandial responses in free-living conditions requires knowing when each meal starts and ends, so that the pre-meal baseline, the eating period, and the subsequent postprandial period can be identified. Self-reported meal times can be burdensome, imprecise, and suffer from poor long-term adherence [6]. This motivates the need for wearables that can operate continuously over long periods, automatically detect eating events, and track the physiological responses that follow. Such a device should also capture more than HR. HR responses provide a non-invasive view of the autonomic adjustments to feeding but reveal little about the accompanying vascular changes. Photoplethysmography (PPG) can address this limitation because, in addition to HR, it captures changes in arterial pulse morphology. Morphological features, such as PPG-derived stiffness and reflection indices, are associated with changes in vascular resistance, arterial compliance, and vasomotor tone [10, 40]. A single wearable that combines PPG with sensors for detecting meal starts and ends is therefore attractive: it could both detect meals and characterize the autonomic and vascular responses that follow, entirely in daily life.

The superficial temporal artery (STA) is an attractive site for postprandial monitoring for both methodological and physiological reasons. Methodologically, it addresses a key challenge in free-living monitoring: determining when a meal starts and ends. The STA runs over the temporalis muscle, which contracts during chewing, so a motion sensor placed at this site can potentially detect the jaw movements of mastication and thereby identify eating events [4, 49]. Physiologically, the STA provides access not only to PPG-based HR monitoring but also to a vascular bed distinct from those in which postprandial vascular changes have previously been characterized. After a meal, splanchnic abdominal vasodilation is offset mainly by an increase in cardiac output, and may be accompanied by reduced blood flow and vasoconstriction in some peripheral beds. For example, Doppler studies have reported reduced blood flow in limb arteries [11] and reduced fingertip skin blood flow [35] after meals. However, skin blood flow at these distal sites is also influenced by thermoregulation and ambient temperature, which can modify or mask meal-related changes [35]. The STA, by contrast, is a branch of the external carotid artery that supplies the scalp. It therefore lies outside the autoregulated cerebral circulation [34], and the scalp lacks the arteriovenous anastomoses that dominate thermoregulatory flow changes in the fingers, which may make it less susceptible to thermoregulatory effects. Because the STA bed differs from both the cerebral and the distal peripheral circulations, its postprandial response may not resemble either. It could show the vasoconstriction reported at distal sites, remain relatively stable, or respond differently altogether. Characterizing this response may therefore provide additional insight into how postprandial hemodynamic adjustments differ across vascular beds.

Here, we used Temple, a wearable device worn over the superficial temporal artery, to examine postprandial cardiovascular responses in 100 healthy adults under free-living conditions. The temple location enables a single device to detect meal onset and offset through chewing-related movements of the temporalis muscle while simultaneously recording PPG signals from the superficial temporal artery. This integration allows postprandial responses to be characterized for each meal without requiring manual logging. Combining the detected meal times with meal composition estimated from photographs, we analyzed the 120 min period after each meal with two aims. First, to assess feasibility, we tested whether postprandial HR responses established in prior studies could be observed from automatically detected meals in daily life. We hypothesized that HR would rise after a meal and then recover toward baseline, and that higher-calorie meals would be associated with a larger and more prolonged rise. We further hypothesized that sodium density and fiber density would be associated with the HR response. As an exploratory analysis, we examined whether the association between meal composition variables and the HR response differed with body mass index (BMI). Second, we explored the vascular response, examining how *A_d_/A_a_*, a PPG morphology marker associated with wave reflection and vascular tone, changed after meals at the temple region, a vascular bed whose postprandial behavior has received little attention. Together, these analyses assess whether a temple-worn wearable can capture cardiac and vascular responses to meals in daily life, a step toward passive monitoring of postprandial cardiovascular health.

## 2 Methods

### 2.1 Device and scope

Temple is a wearable device worn over the superficial temporal artery. It contains a three-axis inertial measurement unit (IMU) positioned over the temporalis muscle sampled at 26 Hz and a green-light PPG sensor sampled at 32 Hz. Data were recorded during participants’ routine use of the device in daily life and analyzed retrospectively (Figure 1A, C).

**Figure 1.**
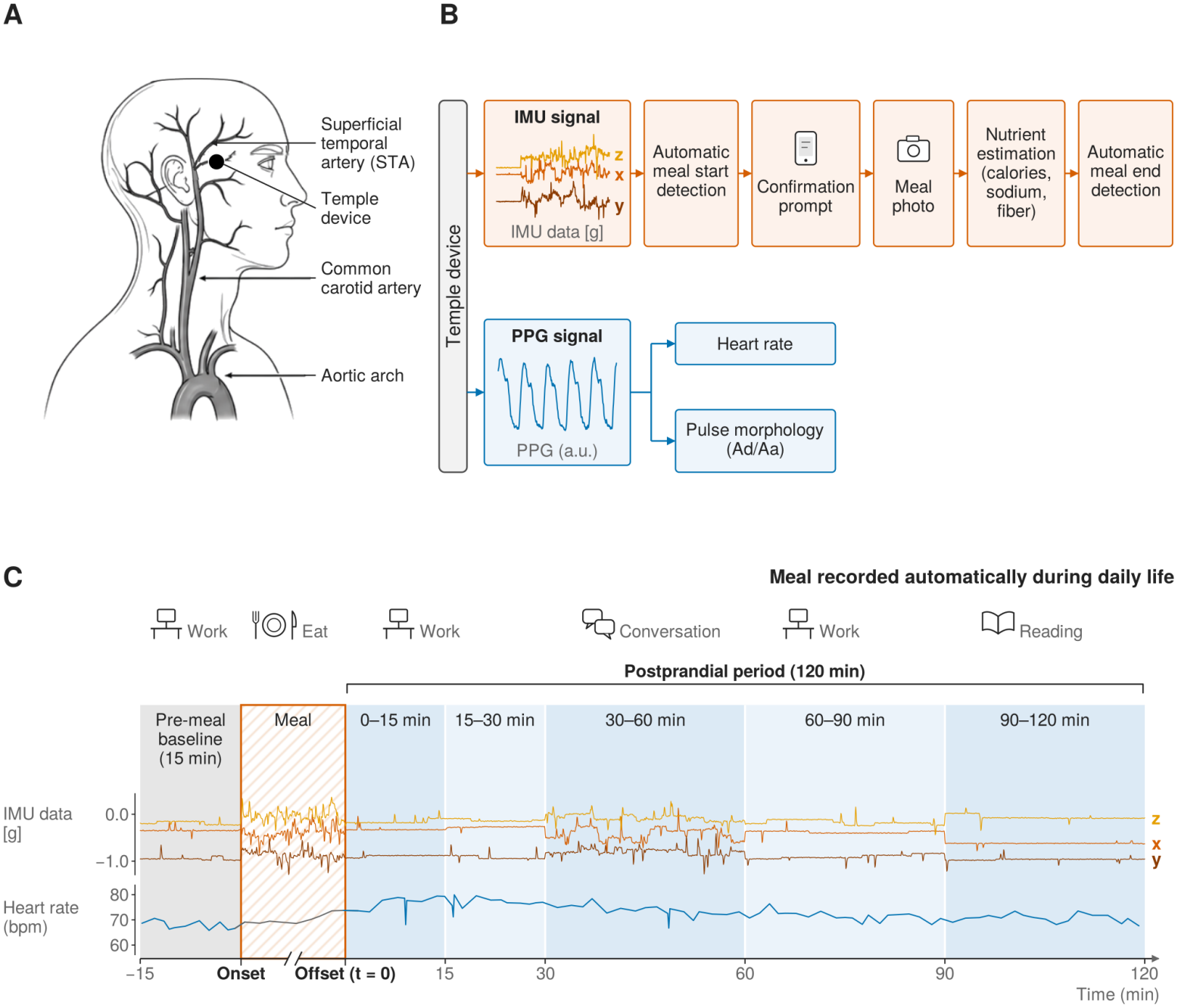
Study setup and data pipeline. **(A)** Device placement. The Temple device is worn at the temple region. **(B)** Data pipeline. The device records two signals. In the inertial measurement unit (IMU) branch (orange), meal onset is detected automatically from chewing-related movement, participants are prompted to confirm the event and photograph the meal, nutrient content (calories, sodium and fiber) is estimated from the photograph, and meal offset is detected automatically from the IMU. In the photoplethysmography (PPG) branch (blue), the PPG signal is used to estimate heart rate and to extract the pulse morphology features such as *A_d_/A_a_*. **(C)** Illustration of an example meal recorded automatically during daily life. IMU signals and heart rate are shown around a detected meal, with the participant’s activities indicated above. The pre-meal baseline (grey) covers the 15 min before meal onset; data recorded during the meal (hatched orange), whose duration varied, were excluded from analyses. Time is measured from meal offset (*t* = 0), and the 120-min postprandial period is divided into five non-overlapping windows (blue): 0–15, 15–30, 30–60, 60–90 and 90–120 min.

As shown in Figure 1B, this study used several outputs generated by the device’s software, together with the raw PPG and accelerometer signals: eating onset and offset, detected automatically from IMU signals; HR, estimated from PPG; and meal nutrient content, estimated from meal photographs. Meal onset, offset, and nutrient content were recorded in a meal log generated in four stages. First, a proprietary meal-detection algorithm identified eating events from IMU data. Second, at each detected onset, participants were prompted to confirm the event and photograph the meal; unconfirmed detections were excluded from the log. Third, a proprietary computer-vision algorithm identified the food items in each photograph and assigned nutrient values to each item. Finally, meal offset was detected from the IMU data.

The accuracy of HR estimation, meal onset and offset detection, and nutrient estimation is evaluated internally against reference measures in studies separate from the present one; these evaluations are essential for establishing full feasibility and will be reported separately.

The present study does not assess the accuracy of individual components. Instead, it examines whether these components, when integrated and operating under free-living conditions, can capture the postprandial HR responses reported in previous studies. It also characterizes meal-related changes in PPG pulse morphology measured at the temple.

### 2.2 Data collection

#### 2.2.1 Meal selection and eligibility criteria

Analyses were conducted on free-living meals recorded by healthy Temple users in their daily settings between 1 April and 30 August 2026. One eligible meal was retained per participant (*N* = 100), so that each observation is independent from each other.

For each meal, PPG signals and PPG-derived HR were extracted from two fixed, non-overlapping periods: a 15-min pre-prandial baseline ending at eating onset, and a 120-min postprandial window beginning at eating end. Eating onset and end were obtained from the meal log generated by automatic eating-episode detection, and meal composition was estimated from photographs of each meal. Data recorded between eating onset and end were excluded from analyses, because mastication can degrade PPG signal quality and the accuracy of HR estimation. The postprandial window was set to 120 min because the largest postprandial changes in cardiac output and HR tend to occur within this period, and after small meals cardiac output returns close to baseline within 120 min [45]. This window is also consistent with previous studies of postprandial changes in central blood pressure, arterial stiffness, cardiac output and HR [2, 18].

Meal timing and nutritional composition were derived from each participant’s meal log. Meal sessions used in our analyses were required to satisfy the following criteria.

##### Meal characteristics

1. **Energy content.** Total energy ≥300 kcal, so that each meal session represents a substantive intake rather than a snack.
2. **Complete nutritional composition.** The meal carried non-null sodium, calories and fiber values.

##### Contextual isolation

– **No contaminating events.** No other logged meal or exercise sessions overlapped the baseline window, the postprandial window, or the 120 min preceding the baseline window. The pre-baseline lookback ensures that the baseline does not itself reflect the lingering physiological effect of an earlier event such as meal or exercise.

##### Wear state and signal quality

– **Continuous valid wear.** Meals were included only if proprietary algorithms confirmed that the participant was awake and the device was worn continuously throughout both the baseline and postprandial windows.
– **Low motion.** Mean accelerometer standard deviation was *<* 0.1 g in both the baseline and the postprandial window. This constraint is intended to exclude ambulation and other physical activity that would itself elevate HR and confound the postprandial HR response, and it additionally suppresses motion artefact in the PPG, supporting accurate HR estimation and PPG morphology analyses.

##### Validation of the motion threshold

To confirm that the 0.1 g ceiling meaningfully separates rest from ambulation, we measured accelerometer activity during 15 tagged light activity sessions from 15 distinct participants with the temple-worn wearable, including slow outdoor walks and indoor walks. Mean accelerometer standard deviation across these sessions was 0.302 *±* 0.122 g, roughly three times the resting ceiling, and no session fell below 0.1 g.

#### 2.2.2 Cohort information

Table 1 summarizes the demographic characteristics of the analytic sample, which comprised 100 meals from 100 participants (one meal per participant) that met the eligibility criteria. Note that amongst the analyzed population, 4 were missing BMI information.

**Table 1:** Characteristics of the analytic sample. One meal session per participant (*N* = 100); all sessions satisfied the eligibility criteria described in Section 2.2. Values are mean *±* SD unless otherwise indicated.

| Variable | *N* | Value |
|---|---|---|
| *Participant characteristics* |  |  |
| Age (years) | 100 | 27.4 ± 4.9 |
| Sex, *N* | 100 |  |
| Male |  | 76 |
| Female |  | 24 |
| BMI (kg/m<sup>2</sup>) | 96 | 23.7 ± 3.5 |
| *Meal characteristics* |  |  |
| Meal type, *N* | 100 |  |
| Breakfast (06:00–10:59) |  | 11 |
| Lunch (11:00–14:59) |  | 58 |
| Dinner (16:00–23:59) |  | 31 |
| Eating duration (min) | 100 | 17.2 ± 14.9 |
| Energy (kcal) | 100 | 713 ± 308 |
| Sodium density (mg/100 kcal) | 100 | 171.1 ± 77.1 |
| Fiber density (g/100 kcal) | 100 | 1.30 ± 0.66 |

### 2.3 Meal composition variables

Three composition variables were analyzed: total energy (kcal), sodium density (mg per 100 kcal) and fiber density (g per 100 kcal). Total energy was taken as the meal’s recorded total calories. The two densities were formed by dividing the nutrient amount by the meal’s total energy and multiplying by 100, following a per-100 kcal food-density definition [47].

Densities rather than raw nutrient amounts were used because raw amounts all scale with the quantity of food consumed and are therefore strongly inter-correlated, which inflates the variance of the coefficient estimates when each nutrient’s contribution is estimated while adjusting for the others. Across the 100 meals, raw amounts were strongly correlated with one another (energy–sodium *r* = 0.72, energy–fiber *r* = 0.54, sodium–fiber *r* = 0.48). Expressing the nutrients as densities attenuated the inter-correlation: (energy–sodium density *r* = 0.05, energy–fiber density *r* = *−* 0.16, sodium–fiber density *r* = 0.29).

### 2.4 Analysis windows

Time was measured from the end of eating (*t* = 0). Each participant’s baseline was the 15 min immediately before eating onset. The first 120 min after the meal were analyzed in two ways (Figure 2). *Non-cumulative* windows divided this period into five consecutive, non-overlapping intervals, [0, 15), [15, 30), [30, 60), [60, 90) and [90, 120) min, to analyze how the response evolves over time. A *cumulative* window, [0, 120) min, was also used to capture overall response.

**Figure 2.**
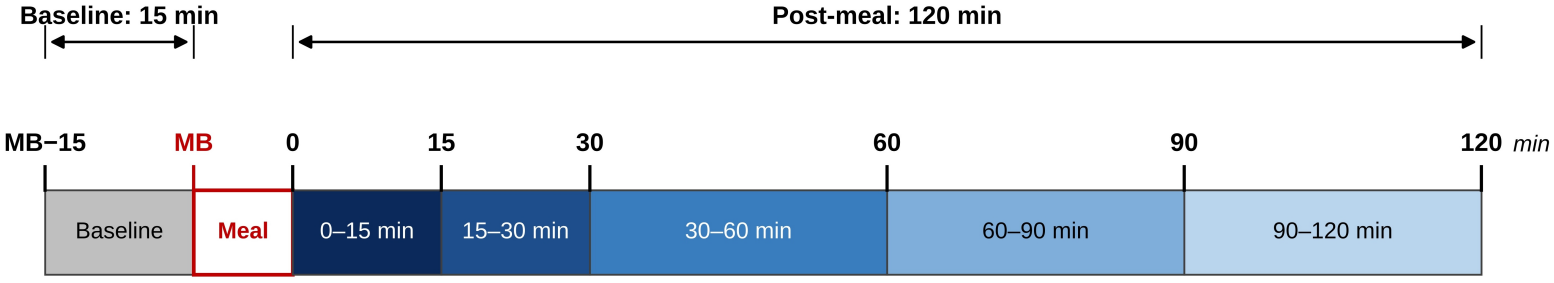
Timeline of baseline and post-meal measurement windows. Times along the top are given relative to the meal. The baseline (grey) covers the 15 min before the meal begins, from MB *−* 15 to MB (MB = meal begins). The meal is shown in red and is not drawn to scale, because its duration varied. Post-meal times are counted from the end of the meal (0 min). The 120 min after the meal is divided into five non-overlapping intervals (blue): 0–15, 15–30, 30–60, 60–90 and 90–120 min. The bracket above the bar shows the cumulative window (0–120 min), starting at 0 min and ending at its upper bound.

### 2.5 Heart rate estimation

Heart rate (HR) was estimated from the PPG signal every 5 s using Temple’s proprietary algorithm. For each participant, HR was summarized as the median of all estimates within each analysis window, yielding seven HR values per participant (*N* = 100), one for each window defined in Section 2.4.

### 2.6 Pulse waveform features

#### 2.6.1 Preprocessing

Pulse-morphology features are more sensitive to signal quality than HR, because they depend on the shape of individual pulses rather than on their timing alone. In addition to the session-level motion criterion described above, we therefore applied a sequence of beat-level quality controls before feature extraction. All analyses used the green PPG channel, sampled at 32 Hz.

The PPG signal was first band-pass filtered between 0.5 and 10 Hz using a fourth-order zero-phase Butterworth filter, and then inverted so that the systolic peak appears as an upward deflection. The 0.5 Hz lower cutoff attenuates slow oscillations such as Mayer waves (*∼*0.1 Hz) and respiratory oscillations (*∼*0.2–0.3 Hz) [21], making fiducial-point detection more robust.

Motion artifacts were then identified at a finer temporal resolution than the session-level criterion. The accelerometer signal was divided into 5-s blocks, and any block whose standard deviation equaled or exceeded 0.05 g (half the threshold applied to whole windows during session selection) was marked as artifactual.

Individual beats were detected on the filtered signal using the pyPPG toolbox [15], and three quality criteria were then applied in sequence. First, beats overlapping an artifactual accelerometer block were discarded. Second, each remaining beat was resampled to 100 points across its onset-to-onset interval, min–max normalized to the range 0–1, and compared with a template; beats whose correlation with the template was below *r* = 0.7 were rejected. Templates were computed as the mean of the normalized beats that passed the motion criterion, separately for the baseline and postprandial periods, pooled across participants. Third, a beat was retained only if both of its immediate neighbors (among all originally detected beats) also passed the first two criteria, so that beats surrounded by corrupted pulses were excluded even if they appeared clean in isolation.

#### 2.6.2 Feature extraction

We extracted the PPG pulse morphology feature *A_d_/A_a_*, computed per beat from the second derivative of the photo-plethysmogram using pyPPG [15]. Within each cardiac cycle, the second derivative exhibits a characteristic sequence of five waves, labeled *a* through *e*: a positive *a* wave and a negative *b* wave in early systole, followed by the *c* and *d* waves in late systole and the *e* wave in early diastole. *A_d_/A_a_* is the amplitude of the *d* wave relative to that of the *a* wave, both measured from the second-derivative baseline. Because the *d* wave is typically negative and the *a* wave positive, the index typically takes negative values. The *d* wave has been related to the late-systolic pressure augmentation produced by waves reflected from the periphery, and *A_d_/A_a_* has therefore been associated with wave reflection and peripheral vascular tone: it increases with vasodilation and decreases with vasoconstriction [40]. Per-beat feature values were pooled together within an analysis window and summarized by their median, giving one value per participant per window — the same aggregation applied to heart rate.

The preprocessing used to extract pulse waveform features was deliberately conservative, so the morphology analysis has a smaller sample than the HR analysis, which included all 100 sessions. This is because morphology features depend on the shape of each pulse rather than only on beat timing. Of these 100 samples, 52 had usable morphology at baseline after applying the artifact and signal-quality criteria, and between 56 and 64 had usable morphology in each postprandial window defined in Section 2.4.

### 2.7 Statistical analyses

#### 2.7.1 Postprandial perturbation and recovery

We first asked whether a meal perturbs HR and pulse morphology at all, and whether the perturbation follows a curved trajectory trend consistent with a rise followed by a return. Each outcome (HR and *A_d_/A_a_*) was modeled across all six timepoints, namely the pre-meal baseline and the five postprandial bins, using a linear mixed-effects model with linear and quadratic time terms and a random intercept for each participant:

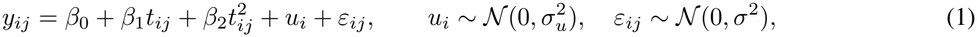

where *y_ij_* is the outcome for participant *i* at timepoint *j, t_ij_* is the mean-centered time in hours, *β*_1_ and *β*_2_ are the linear and quadratic time coefficients, *u_i_* is the participant random intercept, which allows each participant their own baseline level, and *ε_ij_* is the residual error. For each outcome, we report *β*_2_ with its 95% confidence interval and *p* value, together with the number of participants contributing.

To localize the response in time, each postprandial window was then compared with the same participant’s baseline. Because the paired *t* test assumes normally distributed within-participant differences, the differences were tested for normality using the Shapiro–Wilk test (*α* = 0.05); a paired *t* test was used when the differences were consistent with normality, and a Wilcoxon signed-rank test otherwise. Participants missing data from either period were excluded from that comparison only, so *N* varies between windows. For each window, we report the mean change and its standard error, the Cohen’s d effect size *d_z_, N*, and the *p* value. Note that mixed model includes everyone with at least one usable window, while the paired tests need both baseline and that window.

#### 2.7.2 Adjustment for the concurrent HR change

A meal may raise HR as well as altering pulse morphology, and morphology features such as *A_d_/A_a_* may vary mechanically with HR. For *A_d_/A_a_*, the paired comparisons with baseline were therefore repeated after removing the concurrent HR change. For each window, the participants’ changes in *A_d_/A_a_* from baseline were regressed on their changes in HR over the same window:

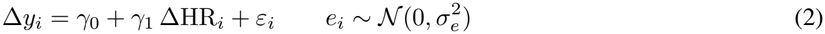

where Δ*y_i_* and ΔHR*_i_* are participant *i*’s changes in *A_d_/A_a_* and HR from baseline, respectively, and *ε_i_* is the residual error. The intercept *γ*_0_ estimates the change in *A_d_/A_a_* that is not explained by the concurrent HR change, that is, the expected change for a participant whose HR did not change. We report *γ*_0_, its 95% confidence intervals and *p* values.

#### 2.7.3 Effects of meal composition

We next asked whether meal composition shapes the time course and the magnitude of the postprandial HR response.

To examine the time course, we extended the recovery model above with a composition variable and its interactions with both time terms:

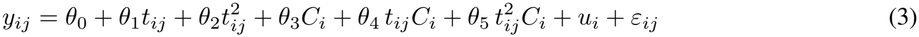

where *C_i_* is the mean-centered composition variable for participant *i*, and all other terms, including the random intercept *u_i_*, are as defined above. The interaction coefficients *θ*_4_ and *θ*_5_ estimate how composition modifies the linear trend and the curvature of the trajectory, respectively. Our sample size may be underpowered to include all three composition variables and their interactions in a single model, so a separate model was fitted for each composition variable. For each model, we report *θ*_4_ and *θ*_5_ with their 95% confidence intervals and *p* values, together with *N*.

To examine the magnitude of the response, all three composition variables were entered into a single model, with the baseline value as a covariate:

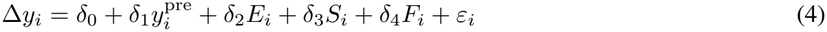

where 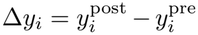 is the change from baseline for participant *i*, 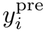 is the baseline value, *E_i_* is total calories (in units of 100 kcal), *S_i_* is sodium density (mg per 100 kcal), *F_i_* is fiber density (g per 100 kcal), and *ε_i_* is the residual error. Because all three composition variables enter the same model, each coefficient estimates the effect of one variable with the other two held constant. This model was fitted separately for each discrete postprandial window and each cumulative window defined in Section 2.4. For each composition variable and window, we report the estimated coefficient with its 95% confidence interval and *p* value, together with *N*.

As a secondary analysis, we compared participants whose meals fell at opposite extremes of each composition variable. For each variable, meals were ranked and divided into tertiles of approximately equal size, and the middle tertile was excluded. For each window, the change from baseline in the highest tertile was compared with that in the lowest tertile, treating the two tertiles as independent groups. Normality of each group’s change scores was assessed with the Shapiro–Wilk test: Welch’s *t* test was used when both groups were consistent with normality, and the Mann–Whitney *U* test otherwise. To complement this between-group contrast, each tertile was also compared with its own pre-meal baseline using the paired procedure described above. For each composition variable and window, we report the group means and standard errors, Cohen’s *d_z_* effect sizes for the paired and unpaired analyses, *p* values, and *N*.

#### 2.7.4 Moderation by BMI

The final question was whether the association between meal composition and the postprandial HR response differs with body mass index (BMI). One model was fitted per composition variable (calories, sodium density, fiber density) at each discrete postprandial bin and each cumulative window, carrying all three composition variables as main effects and a single interaction term:

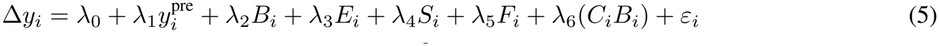

where *B_i_* is BMI, mean-centered and expressed in units of 5 kg/m^2^; *E_i_, S_i_* and *F_i_* are the three composition variables, all mean-centered; *C_i_* is whichever of them is under test in that model; and *λ*_6_ is the moderation effect of interest. Only one interaction is fitted at a time, because relatively small sample size would not provide adequate power to estimate all interactions simultaneously.

For display and interpretation, the same model was refitted with BMI entered as two categorical groups divided at the WHO overweight threshold (BMI *<* 25 versus *≥* 25 kg/m^2^) in place of the continuous term [48]. That model yields each composition variable’s slope within each group.

We report the interaction coefficient with its 95% confidence interval, *N*, *p* values, together with the within-stratum slopes and their 95% confidence intervals.

#### 2.7.5 Multiple comparisons

Analyses were classified as primary or secondary. Primary analyses tested prespecified hypotheses and were not corrected for multiple comparisons. Secondary analyses were considered exploratory and were corrected using the Benjamini–Hochberg procedure, with each composition variable treated as a separate family.

The postprandial HR trajectory model and the comparisons of each postprandial window with baseline were primary. For the analyses of meal composition, the joint regression over the cumulative 0–120 min window and the composition-by-time interaction models were primary. The per-window regressions, including the 90–120 min window used to assess recovery, were secondary and were corrected across the postprandial windows, separately for each composition variable. The tertile analyses were also secondary: the contrasts between the high and low tertiles were corrected across the windows separately for each composition variable, and the within-tertile comparisons with baseline were corrected across the five windows separately for each composition variable and tertile.

All analyses of moderation by BMI were secondary. For each composition variable, the composition-by-BMI interaction over the cumulative 0–120 min window and the five per-window interactions were corrected.

All analyses of the PPG morphology feature *A_d_/A_a_* were secondary. The unadjusted comparisons of each postprandial window with baseline, the HR-adjusted comparisons, and the coefficients on the HR change were each corrected across the analyzed windows.

### 2.8 Ethics approval

The study protocol for data collection received ethical approval from SER-IEC (Approval No. SER-IEC/2026/AP/031; Track No. SER-IEC-NR-2026-025). Written informed consent was obtained from all participants for the use of their data for research purposes.

## 3 Results

### 3.1 Heart rate rises after a meal and approaches baseline by two hours

Mean HR increased significantly by approximately 2.6 bpm during the first 15 min after meal completion. Although HR remained, on average, approximately 2 bpm above baseline thereafter, this difference was not statistically significant. HR subsequently returned to near-baseline levels during the final observation window (Figure 3). A linear mixed-effects model of HR on time confirmed a curved trajectory, with a significant negative quadratic term (*β*_2_ = *−*2.177 bpm h^−2^, 95% CI [*−*3.677*, −*0.678], *p* = 0.004, *N* = 100).

**Figure 3.**
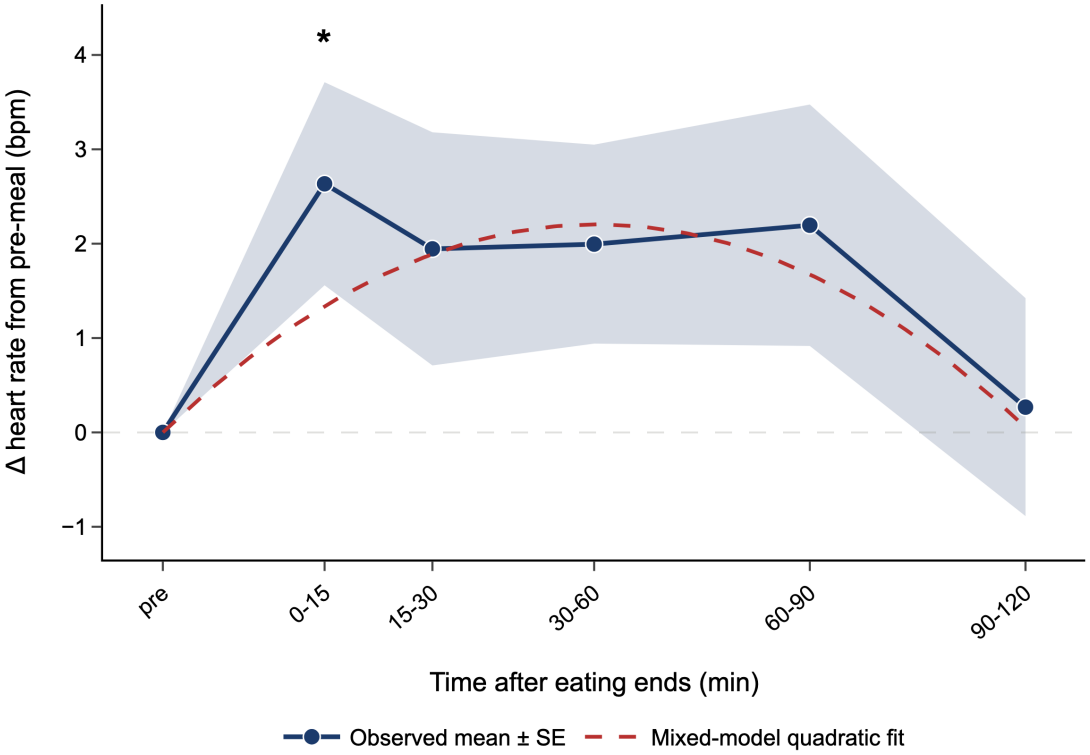
Heart rate rises after a meal and approaches pre-meal baseline by two hours. Change in heart rate from each participant’s own pre-meal baseline across five non-overlapping post-meal windows, timed from the end of eating (*N* = 100 participants; *N* = 97–100 per window after pairwise exclusion of participants missing either side). Points show the across-participant mean and the shaded band *±*1 SE; the pre-meal point is 0 by construction. The dashed curve is the fitted quadratic from a linear mixed-effects model of heart rate on time with a random intercept per participant. The asterisk marks the post-meal window(s) in which heart rate differed significantly from the participants’ own baseline (*p <* 0.05).

Within-participant comparisons of each post-meal window against that participant’s own pre-meal baseline showed that HR was significantly elevated in the first window, 0–15 min after eating ended (Δ = 2.635 *±*1.075 bpm, mean *±*SE; Cohen’s *d_z_* = 0.245, *p* = 0.020, *N* = 100). In the following windows, the mean change remained positive but did not reach significance: 15–30 min (Δ = 1.945 *±*1.235 bpm, *d_z_* = 0.157, *p* = 0.132, *N* = 100), 30–60 min (Δ = 1.995 *±*1.054 bpm, *d_z_* = 0.191, *p* = 0.061, *N* = 98) and 60–90 min (Δ = 2.196 *±*1.279 bpm, *d_z_* = 0.174, *p* = 0.089, *N* = 97). By the final window, 90–120 min, both the mean change and the effect size were the smallest of any post-meal window (Δ = 0.268 *±* 1.155 bpm, *d_z_* = 0.024, *p* = 0.817, *N* = 97).

### 3.2 Meal composition is associated with the magnitude and time course of the postprandial heart-rate rise

We tested whether the postprandial HR trajectory is related to meal composition using three meal parameters: total calories, sodium density (mg per 100 kcal) and fiber density (g per 100 kcal). Extending the trajectory model of Figure 3 with the composition variable and its interactions with both time terms (*N* = 100 for all three models), the curvature interaction was negative and significant for all three (calories: *θ*_5_ = *−* 0.521 bpm h^−2^ per 100 kcal, 95% CI [*−* 1.003*, −* 0.039], *p* = 0.034; sodium density: *θ*_5_ = *−* 0.030 bpm h^−2^ per mg/100 kcal, [*−* 0.049*, −* 0.011], *p* = 0.003; fiber density: *θ*_5_ = *−* 3.554 bpm h^−2^ per g/100 kcal, [*−* 5.798*, −* 1.309], *p* = 0.002). The interaction with the linear time term was positive and significant for calories (*θ*_4_ = 0.497 bpm h^−1^ per 100 kcal, [0.199, 0.795], *p* = 0.001) and sodium density (*θ*_4_ = 0.016 bpm h^−1^ per mg/100 kcal, [0.004, 0.028], *p* = 0.010), but not for fiber density (*θ*_4_ = 0.595 bpm h^−1^ per g/100 kcal, [− 0.793, 1.983], *p* = 0.401). These suggest that the three composition variables might be associated with the shape of the HR trajectory.

The HR change was then regressed on all three meal parameters jointly, adjusted for pre-meal HR. At the cumulative 0–120 min window, the effect of total calories was significant (*δ*_2_ = 0.848 bpm per 100 kcal, 95% CI [0.289, 1.406], *p* = 0.003, *N* = 100). Over the same window the effect of sodium density was weaker but still significant (*δ*_3_ = 0.025 bpm per mg/100 kcal, [0.002, 0.048], *p* = 0.031, *N* = 100), and that of fiber density marginal and non-significant (*δ*_4_ = 2.331 bpm per g/100 kcal, [*−* 0.361, 5.023], *p* = 0.089, *N* = 100). All three slopes were positive, so a higher value of a composition variable corresponded to a larger rise. Resolved window by window, however, the three curves behaved differently. The calorie coefficient was significant in four of the five windows (Figure 4A; 0–15 min: *δ*_2_ = 0.672 [0.083, 1.260], *p_corrected_* = 0.034, *N* = 100; 30–60 min: *δ*_2_ = 0.942 [0.425, 1.459], *p_corrected_* = 0.002, *N* = 98; 60–90 min: *δ*_2_ = 0.896 [0.217, 1.576], *p_corrected_* = 0.025, *N* = 97; 90–120 min: *δ*_2_ = 0.742 [0.084, 1.400], *p_corrected_* = 0.034, *N* = 97), the exception being 15–30 min (*δ*_2_ = 0.512 [*−* 0.138, 1.162], *p_corrected_* = 0.121, *N* = 100). The fiber density effect was significant only in the first window after corrections (Figure 4C; 0–15 min: *δ*_4_ = 4.609 [1.768, 7.449], *p_corrected_* = 0.009, *N* = 100; all later windows *p_corrected_ ≥* 0.05). Sodium density did not reach significance in any individual window (Figure 4B; all *p_corrected_ ≥* 0.05).

**Figure 4.**
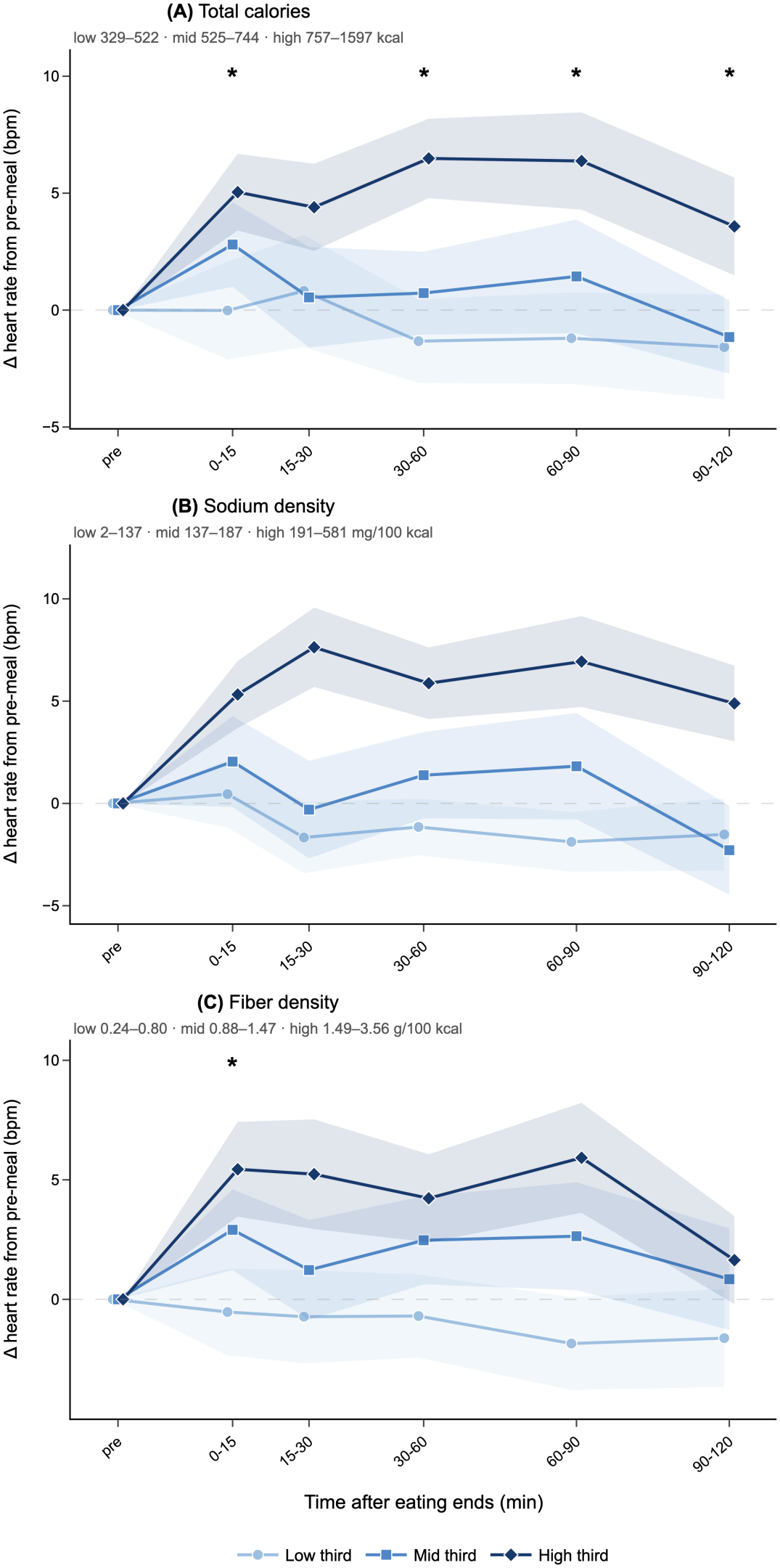
The postprandial heart-rate rise is associated with meal composition. Change in HR from each participant’s own pre-meal baseline across five non-overlapping post-meal windows, timed from the end of eating, with the cohort divided into rank-based tertiles of each composition variable: **(A)** total calories, **(B)** sodium density and **(C)** fiber density. Each legend entry gives that tertile’s observed composition range per tertile (*N* = 31–34). Lines show the mean across participants and the shaded bands show *±* 1 SE; the pre-meal value is 0 by construction. Asterisks mark windows in which that composition variable’s coefficient is significant (*p <* 0.05) in the joint regression of HR change on baseline heart rate and all three composition variables; they do not indicate differences between the plotted tertiles.

In a secondary analysis, we divided participants into tertiles of each composition variable (*N* = 31–34 per group) and compared the high and low tertiles at each window, with the middle tertile excluded. The two outer groups separated most consistently on sodium density, which differed at every window, in contrast to the joint regression, in which sodium density was the weakest predictor at individual windows (low vs high tertile, 0–15 min: Δ = 0.455 *±*1.636 vs Δ = 5.324 *±*1.638 bpm, *d_z_* = 0.514, *p_corrected_* = 0.039, *N* = 33 vs 34; 15–30 min: Δ = *−*1.667 *±*1.741 vs Δ = 7.632 *±*1.939 bpm, *d_z_* = 0.870, *p_corrected_* = 0.003, *N* = 33 vs 34; 30–60 min: Δ = *−* 1.152 *±*1.379 vs Δ = 5.875 *±*1.746 bpm, *d_z_* = 0.786, *p_corrected_* = 0.004, *N* = 33 vs 32; 60–90 min: Δ = *−*1.879 *±*1.456 vs Δ = 6.935 *±*2.226 bpm, *d_z_* = 0.839, *p_corrected_* = 0.004, *N* = 33 vs 31; 90–120 min: Δ = *−* 1.515*±* 1.766 vs Δ = 4.887 *±* 1.847 bpm, *d_z_* = 0.627, *p_corrected_* = 0.019, *N* = 33 vs 31). Calories separated the tertiles at 30–60 and 60–90 min (30–60 min: Δ = *−*1.328 *±*1.781 vs Δ = 6.485 *±*1.696 bpm, *d_z_* = 0.788, *p_corrected_* = 0.012, *N* = 32 vs 33; 60–90 min: Δ = *−*1.203 *±*1.950 vs Δ = 6.375 *±*2.079 bpm, *d_z_* = 0.665, *p_corrected_* = 0.036, *N* = 32 vs 32), and fiber density did not show between tertile significance at the analyzed windows (*p_corrected_ ≥* 0.05), after corrections for multiple comparisons.

Within-group comparisons against each group’s own pre-meal baseline showed that the high tertiles rose above their own baselines, whereas changes in the low tertiles were small and not significant in any window of any composition variable (*p_corrected_ ≥* 0.05). For calories, the high tertile rose at 0–15 min (Δ = 5.044 *±* 1.636 bpm, *d_z_* = 0.529, *p_corrected_* = 0.007, *N* = 34), 30–60 min (Δ = 6.485 *±* 1.696 bpm, *d_z_* = 0.665, *p_corrected_* = 0.003, *N* = 33) and 60–90 min (Δ = 6.375 *±* 2.079 bpm, *d_z_* = 0.542, *p_corrected_* = 0.007, *N* = 32), and was no longer significantly elevated by 90–120 min (Δ = 3.578 *±* 2.090 bpm, *d_z_* = 0.303, *p_corrected_* = 0.097, *N* = 32), while changes in the low tertile were small and not significant (*p_corrected_ ≥* 0.05). For sodium density, the high tertile was significantly above its own baseline in every window (0–15 min: Δ = 5.324 *±* 1.638 bpm, *d_z_* = 0.557, *p_corrected_* = 0.004, *N* = 34; 15–30 min: Δ = 7.632 *±* 1.939 bpm, *d_z_* = 0.675, *p_corrected_* = 0.002, *N* = 34; 30–60 min: Δ = 5.875 *±* 1.746 bpm, *d_z_* = 0.595, *p_corrected_* = 0.004, *N* = 32; 60–90 min: Δ = 6.935 *±* 2.226 bpm, *d_z_* = 0.560, *p_corrected_* = 0.004, *N* = 31; 90–120 min: Δ = 4.887 *±* 1.847 bpm, *d_z_* = 0.475, *p_corrected_* = 0.013, *N* = 31), and the low tertile in none (*p_corrected_ ≥* 0.05). For fiber density, the high tertile rose at 0–15 min (Δ = 5.441 *±* 1.982 bpm, *d_z_* = 0.471, *p_corrected_* = 0.037, *N* = 34), 30–60 min (Δ = 4.227 *±* 1.840 bpm, *d_z_* = 0.400, *p_corrected_* = 0.047, *N* = 33) and 60–90 min (Δ = 5.922 *±* 2.299 bpm, *d_z_* = 0.455, *p_corrected_* = 0.037, *N* = 32), and was no longer elevated by 90–120 min (Δ = 1.641 *±* 1.840 bpm, *d_z_* = 0.158, *p_corrected_* = 0.379, *N* = 32), with no significant change in the low tertile (*p_corrected_ ≥* 0.05).

### 3.3 Heart rate two hours after eating is associated with meal calories and sodium density

At the 90–120 min window — the end of the postprandial period studied— the high tertile of each composition variable lay above the mid and low tertiles (Figure 4), suggesting that participants’ HR might still be perturbed at that timestamp and take longer to return to baseline.

In the joint model at this final window, total calories had a significant effect (*δ*_2_ = 0.742 bpm per 100 kcal, 95% CI [0.084, 1.400], *p_corrected_* = 0.034, *N* = 97), suggesting that calories positively predicted the postprandial HR change with the other composition variables and baseline HR held constant. Neither sodium density (*δ*_3_ = 0.016 bpm per mg/100 kcal, [*−*0.012, 0.043], *p_corrected_* = 0.328, *N* = 97) nor fiber density (*δ*_4_ = 0.739 bpm per g/100 kcal, [*−*2.454, 3.933], *p_corrected_* = 0.647, *N* = 97) was significantly associated with the change.

In the secondary tertile comparison, calories did not significantly separate the high and low tertiles at this window (Δ = *−*1.578 *±* 2.239 vs Δ = 3.578 *±* 2.090 bpm, *d_z_* = 0.421, *p_corrected_* = 0.127, *N* = 32 vs 32), despite the significant continuous association. The separation was significant for sodium density (Δ = *−*1.515 *±* 1.766 vs Δ = 4.887 *±* 1.847 bpm, *d_z_* = 0.627, *p_corrected_* = 0.019, *N* = 33 vs 31), and not for fiber density (Δ = *−*1.621*±* 2.045 vs Δ = 1.641 *±* 1.840 bpm, *d_z_* = 0.294, *p_corrected_* = 0.318, *N* = 33 vs 32). Thus, at two hours, calories were associated with the HR change in the continuous model, whereas sodium density separated the extreme tertiles but was not significant as a continuous predictor.

Paired comparisons within each tertile showed a similar pattern. For calories, high tertile was not significantly above its own baseline at 90–120 min (Δ = 3.578 *±* 2.090 bpm, *d_z_* = 0.303, *p_corrected_* = 0.097, *N* = 32; mid: Δ = *−*1.152 *±*1.566 bpm, *d_z_* = *−*0.128, *p_corrected_* = 0.801, *N* = 33; low: Δ = *−*1.578 *±*2.239 bpm, *d_z_* = *−*0.125, *p_corrected_* = 0.994, *N* = 32), although its mean change and effect size were the largest of the three tertiles. For sodium density, the high tertile remained significantly above its own baseline (Δ = 4.887 *±* 1.847 bpm, *d_z_* = 0.475, *p_corrected_* = 0.013, *N* = 31), while the mid and low tertiles did not (mid: Δ = *−*2.288 *±* 2.170 bpm, *d_z_* = *−*0.184, *p_corrected_* = 0.648, *N* = 33; low: Δ = *−*1.515 *±* 1.766 bpm, *d_z_* = *−*0.149, *p_corrected_* = 0.513, *N* = 33). For fiber density, no tertile differed from baseline and all effect sizes were small (high: Δ = 1.641 *±* 1.840 bpm, *d_z_* = 0.158, *p_corrected_* = 0.379, *N* = 32; mid: Δ = 0.844 *±* 2.120 bpm, *d_z_* = 0.070, *p_corrected_* = 0.693, *N* = 32; low: Δ = *−*1.621 *±* 2.045 bpm, *d_z_* = *−*0.138, *p_corrected_* = 0.771, *N* = 33).

### 3.4 The HR response to calories is larger in participants with higher BMI

The effect of BMI on the postprandial HR response to meal composition was examined by regressing the HR change from baseline on baseline HR, BMI, all three composition variables, and the interaction between BMI and one composition variable, with a separate model fitted for each composition variable. Over the cumulative 0–120 min window, the calories *×* BMI interaction was significantly positive (*λ*_6_ = 1.080 bpm per 100 kcal per 5 kg/m^2^, 95% CI [0.282, 1.878], *p_corrected_* = 0.027, *N* = 96). Therefore, the HR response to a given caloric load was larger in participants with higher BMI and reached significance in three of the five individual windows (0–15 min: *λ*_6_ = 0.923 [0.069, 1.777], *p_corrected_* = 0.041, *N* = 96; 15–30 min: *λ*_6_ = 1.351 [0.415, 2.287], *p_corrected_* = 0.027, *N* = 96; 60–90 min: *λ*_6_ = 1.181 [0.218, 2.144], *p_corrected_* = 0.027, *N* = 93) and not in the remaining two (30–60 min: *λ*_6_ = 0.658 [*−* 0.108, 1.424], *p_corrected_* = 0.091, *N* = 94; 90–120 min: *λ*_6_ = 0.858 [*−* 0.034, 1.750], *p_corrected_* = 0.071, *N* = 93). Over the same cumulative window, the interaction was marginal and not significant for sodium density (*λ*_6_ = 0.040 bpm per mg/100 kcal per 5 kg/m^2^, [*−* 0.001, 0.082], *p_corrected_* = 0.058, *N* = 96) and not significant for fiber density (*λ*_6_ = 2.334 bpm per g/100 kcal per 5 kg/m^2^, [*−* 1.716, 6.384], *p_corrected_* = 0.255, *N* = 96).

For illustration, Figure 5 shows the same moderation with BMI split at the WHO overweight threshold (*<* 25 versus *≥* 25 kg/m^2^), giving each group its own slope. With baseline heart rate and the other two composition variables held constant, the slope on calories was more than four times as steep in participants with BMI *≥* 25 than in those with BMI *<* 25 (panel A; BMI *≥* 25: 2.348 bpm per 100 kcal, [1.113, 3.582]; BMI *<* 25: 0.525, [*−*0.098, 1.148]). The same ordering was seen descriptively for sodium density (panel B; BMI *≥* 25: 0.072 bpm per mg/100 kcal, [0.017, 0.127]; BMI *<* 25: 0.017, [*−*0.008, 0.043]) and fiber density (panel C; BMI *≥* 25: 3.958 bpm per g/100 kcal, [0.098, 7.817]; BMI *<* 25: 0.408, [*−*3.429, 4.246]), although their interactions with BMI were not significant.

**Figure 5.**
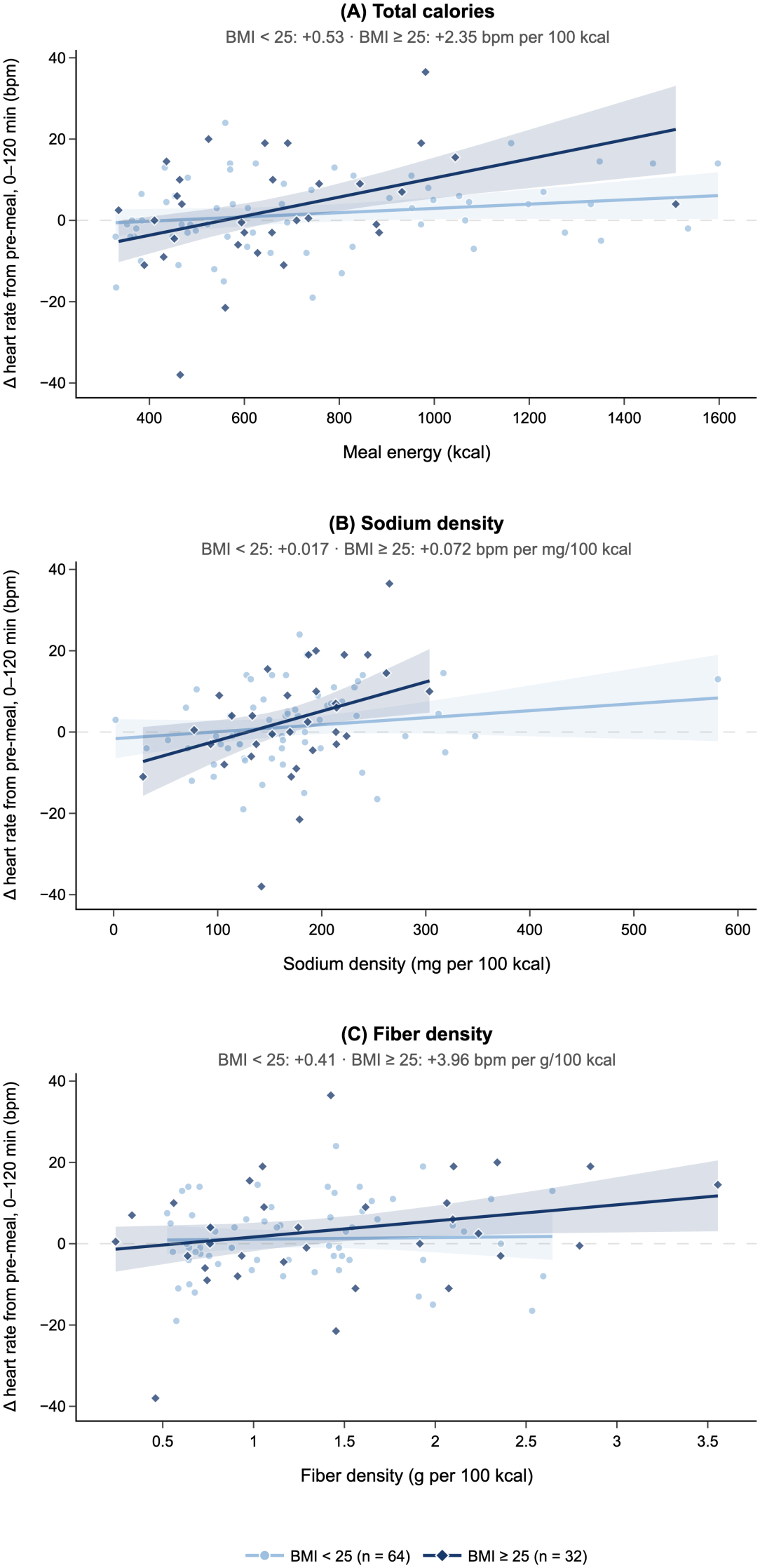
The heart-rate response to meal calories is steeper in participants with higher BMI. Change in heart rate from each participant’s own pre-meal baseline, taken as median over the 0–120 min post-meal window, plotted against **(A)** calories, **(B)** sodium density and **(C)** fiber density. Each point is one participant (*N* = 96 with a recorded BMI: 64 at BMI *<* 25, 32 at BMI *≥* 25), split at the WHO overweight threshold and shown in light circles (BMI *<* 25) and dark diamonds (BMI *≥* 25). Lines are fitted values from an ordinary least-squares model estimated over both BMI groups, in which the slope on the composition variable is allowed to differ between groups and the heart-rate change is adjusted for pre-meal heart rate and the other two composition variables. Each slope is given above its panel.

### 3.5 PPG pulse-waveform changes consistent with vasodilation occur after a meal, largely independent of heart rate

The PPG morphology feature *A_d_/A_a_*, associated with wave reflection and vascular tone, rose after the meal and was still elevated two hours later (Figure 6). In a linear mixed-effects model of *A_d_/A_a_* on time with a random intercept per participant, the quadratic term was negative (*β*_2_ = *−*0.086 a.u. h^−2^, [−0.124, −0.048], *p* < 0.001, *N* = 76). Unlike

**Figure 6.**
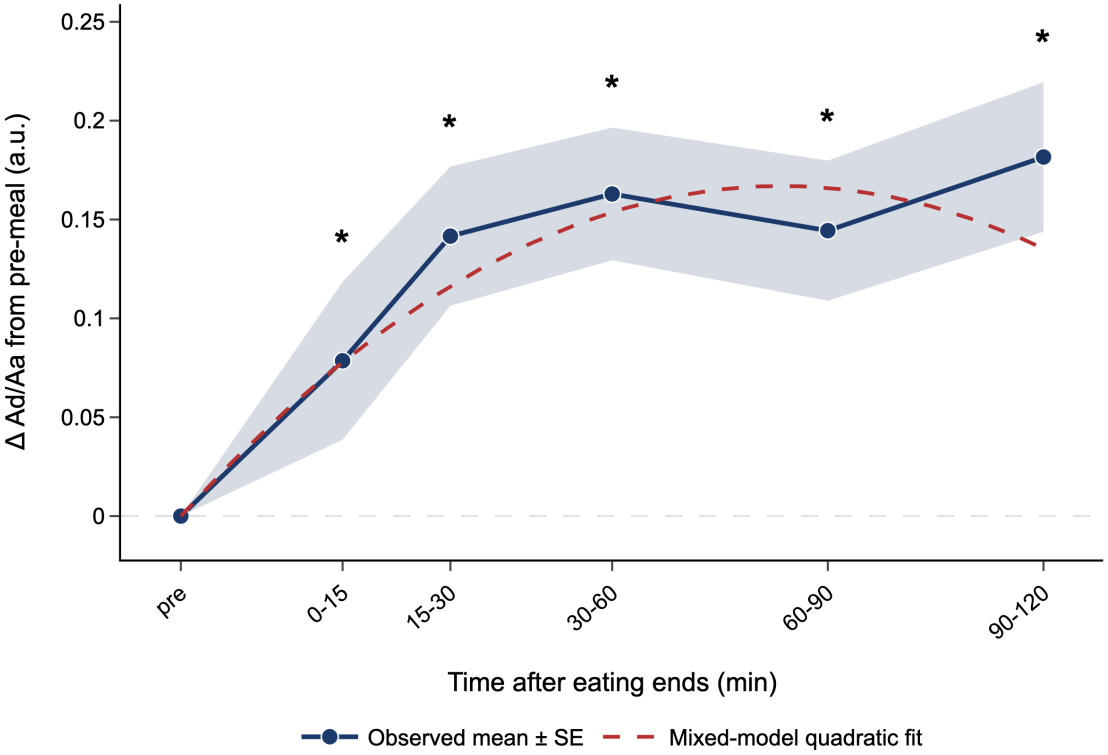
PPG pulse-waveform changes consistent with vasodilation occur after a meal and remain shifted at two hours. Change from each participant’s pre-meal baseline in the green-channel morphology feature *A_d_/A_a_*, which has been associated with wave reflection and vascular tone, across five non-overlapping post-meal windows timed from the end of eating (*N* = 40–44 per window; participants missing either the baseline or the window value were excluded pairwise). Points show the across-participant mean and the shaded band *±*1 SE. The dashed curve is the fitted quadratic from a linear mixed-effects model of *A_d_/A_a_* on time with a random intercept per participant (*N* = 76). The negative curvature indicates a rise that decelerates rather than resolves; note that the observed mean change is largest in the final window. The asterisk marks the post-meal window(s) in which *A_d_/A_a_* differed significantly from the participants’ own baseline (*p <* 0.05).

HR, however, *A_d_/A_a_* did not return toward baseline: the largest mean change occurred in the final window. Together, these suggest a rise that decelerated but did not return to baseline within the two-hour period.

Paired comparisons of each participant’s *A_d_/A_a_* against their own pre-meal baseline locate the elevation in every analyzed window. *A_d_/A_a_* was elevated within 15 min of the meal (Δ = 0.079 *±* 0.040, *d_z_* = 0.311, *p_corrected_* = 0.016, *N* = 40) and remained so throughout: 15–30 min (Δ = 0.142 *±* 0.035, *d_z_* = 0.636, *p_corrected_ <* 0.001, *N* = 40), 30–60 min (Δ = 0.163 *±*0.034, *d_z_* = 0.732, *p_corrected_ <* 0.001, *N* = 44), 60–90 min (Δ = 0.144 *±*0.035, *d_z_* = 0.636, *p_corrected_ <* 0.001, *N* = 41) and 90–120 min (Δ = 0.182 *±* 0.038, *d_z_* = 0.724, *p_corrected_ <* 0.001, *N* = 44).

Because a meal also raises heart rate, and pulse-morphology features can move with heart rate changes too, each window was also tested with the participant’s concurrent heart-rate change regressed out. The adjustment left the estimates close to their unadjusted values, and the elevation remained significant in four of the five windows (15– 30 min: *γ*_0_ = 0.138, 95% CI [0.066, 0.211], *p_corrected_ <* 0.001, *N* = 40; 30–60 min: *γ*_0_ = 0.142, [0.071, 0.214], *p_corrected_ <* 0.001, *N* = 44; 60–90 min: *γ*_0_ = 0.142, [0.066, 0.218], *p_corrected_ <* 0.001, *N* = 41; 90–120 min: *γ*_0_ = 0.177, [0.104, 0.249], *p_corrected_ <* 0.001, *N* = 44), falling to marginal only in the first (0–15 min: *γ*_0_ = 0.078, [*−* 0.003, 0.160], *p_corrected_* = 0.060, *N* = 40). The coefficient on the concurrent heart-rate change in the regression model, *γ*_1_, was close to zero in every window and did not differ significantly from zero in the analyzed windows (*p_corrected_ ≥* 0.05).

## 4 Discussion

In this study, we used a temple-worn photoplethysmography (PPG) wearable with automatic meal detection to characterize postprandial cardiovascular responses in 100 participants under free-living conditions. To test for feasibility, we asked whether the device could recover, in daily life, the postprandial HR response established in prior studies. The device recovered the expected response: HR followed a rise and recovery after a meal. It was significantly elevated during the first 15 min after meal offset, remained modestly above baseline on average up to 90 min, and returned to near baseline by 90–120 min. Higher-calorie meals were associated with a larger HR rise over the two hours after eating, and this association held across most individual postprandial windows and was still present at 90–120 min, indicating a rise that was both larger and more prolonged. In addition to calories, sodium density and fiber density were also associated with the shape of the HR trajectory. Sodium density was associated with a larger rise over the full two hours, although more weakly than calories and in no individual window. In secondary analyses, however, participants eating the most and least sodium-dense meals differed throughout the postprandial period, including at two hours. Fiber density was not associated with the overall rise, and its association was limited to the first 15 min. Participants in the highest tertiles of calories, fiber and sodium density showed HR increases above their own baseline for much of the postprandial period, whereas those in the lowest tertiles did not. In exploratory analyses, the association between caloric content and the HR rise was stronger in participants with higher BMI. Finally, we characterized the postprandial vascular response at the temple, a site that, to our knowledge, has not been previously investigated in this context. The PPG morphology index *A_d_/A_a_* increased after the meal and remained elevated at two hours rather than returning to baseline, and this change was largely unaffected by adjustment for the concurrent HR change, a pattern consistent with reduced arterial wave reflection [40].

The HR response observed here matches that described in prior studies. Postprandial HR rises within the first 30 min after a meal and then declines [23, 18], and the magnitude and duration of the rise scale with meal calories [37, 1, 45]. Sodium density was associated with the HR rise over the two hours, although less than that of calories. In secondary analyses, participants eating the most and least sodium-dense meals differed in every postprandial window, suggesting that very sodium-dense meals may produce a more prolonged HR response. Fiber density showed the least expected pattern. Viscous fiber slows gastric emptying and glucose absorption [20, 36], and adding guar to an intraduodenal glucose infusion attenuated the rise in HR during the first 30 min [32]. Fiber might therefore be expected to produce a smaller, later HR response. Instead, fiber density was not associated with the overall rise, and in secondary analyses higher fiber density was associated with a larger rise in the first 15 min. One explanation is that adjusting for total energy does not account for how that energy is distributed among carbohydrate, fat and protein, which were not modelled here. Meals high in fiber per calorie, such as those based on grains, legumes, fruit and vegetables, tend to be rich in carbohydrate and low in fat, and carbohydrate-rich meals elicit a larger HR increase than fat-rich meals of equal energy [17].

After the meal, *A_d_/A_a_* at the temple became less negative and remained shifted at two hours, and this shift persisted after adjustment for each participant’s concurrent HR change. No such sustained shift was seen in a control window spanning the two hours before the meal. A shallower d-wave indicates a smaller late-systolic component of the pulse, and hence less of the pressure wave returning from the periphery during systole: vasoconstrictors deepen the d-wave relative to the a-wave, whereas the vasodilator nitroglycerin makes it shallower [40]. Therefore, the temple observation is consistent with reduced arterial wave reflection and vasodilation after a meal. Although augmentation indices also decrease as HR increases [46], the persistence of the *A_d_/A_a_* shift after HR adjustment suggests that the change is not explained solely by the postprandial rise in HR, in line with the HR-independent fall in augmentation index reported after a mixed meal [41]. A plausible explanation for this observation is vasodilation downstream of the aorta, chiefly in the gut, where artery flow rises after a meal alongside an increase in cardiac output [12, 38]. As the splanchnic vessels dilate, less of each pressure wave is reflected back up the aorta, so the central pressure waveform might carry a smaller late-systolic component that the carotid circulation could transmit to the superficial temporal artery. Two features of the temple may favor detection of such a central change. First, the superficial temporal artery is a terminal branch of the external carotid artery, so the pulse reaches it by a short, direct path from the aortic arch and may retain changes in central pulse morphology; by contrast, the pulse reaching the finger is reshaped as it travels along the arm [33]. Second, the skin of the forehead, adjacent to the temple, shows little sympathetic vasoconstrictor activity [31], so local vasomotor changes may add little to the transmitted signal. Finger vessels, by contrast, respond readily to sympathetic activity and temperature: during hypothermia-induced vasoconstriction, the response of forehead and ear oximeters was unaffected, whereas that of finger oximeters slowed markedly [29], and during a cold pressor test, PPG features changed most strongly at the finger, with one feature changing in opposite directions at the finger and the head [14]. This difference matters after a meal, because the limbs show their own, opposite response to eating. As part of the compensation for splanchnic vasodilation, limb vessels constrict, and postprandial blood flow falls in the limb arteries [11, 39] and in fingertip skin [35]. A finger or wrist sensor would therefore be expected to capture mainly this local vasoconstriction, whereas the temple may instead reflect the central reduction in wave reflection. Simultaneous recordings at the temple, finger and wrist would be needed to test this.

This study has several limitations. First, several factors limit the evidence for feasibility and for the associations between postprandial responses and meal composition. Most importantly, recovering established postprandial HR trends at the population level shows that the integrated system behaves as expected, but it does not establish the accuracy of its individual components: meal onset and offset detection, image-based nutrient estimation and PPG-derived HR estimation. The accuracy of each component has been evaluated in separate internal studies, which are outside the scope of this work and have not yet been published. These validations are a prerequisite for full feasibility, and the present findings should be interpreted as conditional on them. In addition, only meals that were automatically detected and then confirmed by participants were analyzed; meals the algorithm failed to detect were not captured, and if detection performance differs by meal type or duration, the analyzed meals may not be representative of all meals eaten.

The free-living design that the temple-worn device makes possible, with meals and the responses that follow them recorded automatically, also comes at the cost of experimental control. Meals varied in size, composition and volume, and posture, time of day, social context, conversation, caffeine and alcohol intake, and ambient temperature were not controlled. Although the motion criterion excluded physical activity, it could not exclude other influences on HR such as caffeine, and it restricts the findings to sedentary postprandial periods. In addition, meal composition was not assigned, and composition was estimated from a single photograph by a computer-vision algorithm, and image-based estimation can be error-prone. Ingredients such as salt in sauces and cooking oil may be poorly captured, so sodium and fiber are likely estimated less accurately than total energy. Food left uneaten was not accounted for because the photograph was taken at meal onset, beverages, including caffeinated, alcoholic and sugary drinks, were likely incompletely recorded, and macronutrient content (carbohydrate, fat and protein) was not modeled. The 120-min window may also have been too short for the largest meals, as HR in the highest calorie and sodium tertiles had not fully returned to baseline by two hours (Figure 4), so the full trajectory after the largest meals was not observed. Finally, with one meal per participant, the results do not show whether the same relationships hold within individuals across repeated meals, which is required before the device can be used to interpret an individual’s responses.

Second, the findings were obtained in a relatively small and homogeneous sample and may not generalize beyond it. The cohort consisted of 100 self-reported healthy adults who were young (27.4 *±* 4.9 years), predominantly male (76%) and mostly of normal weight (BMI 23.7 *±* 3.5 kg/m^2^), so the findings may not extend to older adults or clinical populations, in whom tracking of postprandial responses may be most valuable. With only 32 participants with a BMI of 25 kg/m^2^ or above, the BMI moderation analysis was also limited. Participants were users of a consumer wearable and may be more health-conscious than the general population, and characteristics that can influence HR and PPG responses, including ethnicity, skin tone, fitness, medication use, smoking and medical history, were not recorded, so the diversity of the sample on these characteristics is unknown. Larger and more diverse samples are needed to establish how generalizable these results are.

Third, the PPG morphology analyses and *A_d_/A_a_* findings have specific limitations. Because recordings were made under free-living rather than controlled laboratory conditions, the PPG signal was more susceptible to artifacts, and the strict quality control this required left only 40–44 participants with usable data at both baseline and each window. The PPG was sampled at 32 Hz, which is low for second-derivative features such as *A_d_/A_a_*, and higher sampling rates would allow more precise morphology analyses.

Future work should address these limitations. First, it might be valuable to test whether the findings hold under controlled laboratory conditions with the same temple-worn device. Studies in which participants rest in a controlled environment and eat meals of known composition, prepared to vary energy, sodium or fiber independently while volume and temperature are held constant, would establish whether the associations observed here can be replicated. This would also remove the inaccuracies in nutrient estimation. Other factors that can influence the postprandial response, including posture, time of day, ambient temperature, and caffeine and alcohol intake, should also be standardized, so that any change in HR or pulse morphology can be attributed to the meal itself. Second, such conditions would also allow higher-quality recordings of PPG morphology at higher sampling rates. Simultaneous recordings at the temple, finger and wrist would show how postprandial vascular responses differ between sites and test whether the *A_d_/A_a_* findings can be replicated. Third, larger and more diverse cohorts, including older adults, people with overweight or obesity, and clinical populations such as people with coronary artery disease, diabetes or autonomic dysfunction, would test whether these findings extend beyond young, healthy adults, and would include enough participants to examine moderation by BMI, age and sex with adequate power. Longer recording windows, extending beyond two hours, would capture the full recovery of the HR response after the largest meals. Finally, recording repeated meals from each participant would allow within-person analyses, testing whether the relationships between meal composition and the postprandial response observed across individuals also hold within individuals, how consistent each person’s response is from meal to meal, and whether individual response profiles are stable enough to support personalized feedback on the cardiovascular impact of meals.

In summary, we examined postprandial cardiovascular responses in 100 participants in daily life using a single wearable at the temple, a location that permits both automatic meal detection from chewing-related movement and PPG recording from the superficial temporal artery. The HR response agreed with earlier studies: it increased after eating and was close to baseline by two hours, with higher-calorie meals producing a greater and longer-lasting rise. After the most sodium-dense meals, the rise persisted to the end of the two-hour period, whereas fiber density was linked to a larger rise only in the first 15 min. Pulse morphology at the temple also shifted after eating and stayed shifted for two hours. This change was not accounted for by the accompanying rise in HR, pointed to reduced arterial wave reflection, and may contrast with the vasoconstriction described at limb sites after meals. Because the expected HR response and its association with meal energy emerged from meals detected automatically by the device, these results potentially support the feasibility of using a single temple-worn device both to identify eating episodes and to measure the cardiovascular responses that follow in daily life. Full feasibility, however, depends on the accuracy of the device’s individual components, which are evaluated separately and independently from this study. With such validation, together with confirmation in controlled settings, in larger populations and within individuals across repeated meals, these measurements may eventually contribute to a meal-level index of cardiovascular impact, helping individuals understand the cardiovascular effects of their meals.

## References

[1] Eamon Abdullah, Jordan B Lee, Karambir Notay, and Philip J Millar. Postprandial superior mesenteric artery blood flow is related to changes in peripheral pulse wave harmonics and heart rate: implications for wearable technology? Journal of Applied Physiology, 131(2):681–688, 2021.

[2] Kiran DK Ahuja, Iain K Robertson, and Madeleine J Ball. Acute effects of food on postprandial blood pressure and measures of arterial stiffness in healthy humans. The American journal of clinical nutrition, 90(2):298–303, 2009.

[3] R. R. Baliga, S. D. Rosen, P. G. Camici, and J. S. Kooner. Regional myocardial blood flow redistribution as a cause of postprandial angina pectoris. Circulation, 97:1144–1149, 1998.

[4] Abdelkareem Bedri, Richard Li, Malcolm Haynes, Raj Prateek Kosaraju, Ishaan Grover, Temiloluwa Prioleau, Min Yan Beh, Mayank Goel, Thad Starner, and Gregory Abowd. Earbit: using wearable sensors to detect eating episodes in unconstrained environments. Proceedings of the ACM on interactive, mobile, wearable and ubiquitous technologies, 1(3):1–20, 2017.

[5] Rubin Berlinerblau and Jacob Shani. Postprandial angina pectoris: clinical and angiographic correlations. Journal of the American College of Cardiology, 23(3):627–629, 1994.

[6] Lora E Burke, Jing Wang, and Mary Ann Sevick. Self-monitoring in weight loss: a systematic review of the literature. Journal of the American Dietetic Association, 111(1):92–102, 2011.

[7] W. Y. Chung, D. W. Sohn, Y. J. Kim, et al. Absence of postprandial surge in coronary blood flow distal to significant stenosis: a possible mechanism of postprandial angina. Journal of the American College of Cardiology, 40(11):1976–1983, 2002.

[8] K. M. Dickinson, P. M. Clifton, L. M. Burrell, P. H. R. Barrett, and J. B. Keogh. Postprandial effects of a high salt meal on serum sodium, arterial stiffness, markers of nitric oxide production and markers of endothelial function. Atherosclerosis, 232(1):211–216, 2014.

[9] Kacie M Dickinson, Peter M Clifton, and Jennifer B Keogh. Endothelial function is impaired after a high-salt meal in healthy subjects. The American journal of clinical nutrition, 93(3):500–505, 2011.

[10] Mohamed Elgendi. On the analysis of fingertip photoplethysmogram signals. Current cardiology reviews, 8(1):14–25, 2012.

[11] Masako Yamaoka Endo, Chizuko Fujihara, Akira Miura, Hideaki Kashima, and Yoshiyuki Fukuba. Effects of meal ingestion on blood pressure and regional hemodynamic responses after exercise. Journal of Applied Physiology, 120(11):1343–1348, 2016.

[12] M. Eriksen and B. A. Waaler. Priority of blood flow to splanchnic organs in humans during pre- and post-meal exercise. Acta Physiologica Scandinavica, 150(4):363–372, 1994.

[13] Jan Fagius and Christian Berne. Increase in muscle nerve sympathetic activity in humans after food intake. Clinical Science, 86(2):159–167, 1994.

[14] Vincent Fleischhauer, Jan Bruhn, Stefan Rasche, and Sebastian Zaunseder. Photoplethysmography upon cold stress—impact of measurement site and acquisition mode. Frontiers in Physiology, 14:1127624, 2023.

[15] Márton Á Goda, Peter H. Charlton, and Joachim A. Behar. pyPPG: a python toolbox for comprehensive photoplethysmography signal analysis. Physiological Measurement, 45(4):045001, 2024.

[16] D. Neil Granger, Lena Holm, and Peter Kvietys. The gastrointestinal circulation: physiology and pathophysiology. Comprehensive Physiology, 5(3):1541–1583, 2015.

[17] D Heseltine, JF Potter, G Hartley, IA Macdonald, and OFW James. Blood pressure, heart rate and neuroendocrine responses to a high carbohydrate and a high fat meal in healthy young subjects. Clinical Science, 79(5):517–522, 1990.

[18] Joanna Hlebowicz, Sandra Lindstedt, Ola Björgell, and Magnus Dencker. Relationship between postprandial changes in cardiac left ventricular function, glucose and insulin concentrations, gastric emptying, and satiety in healthy subjects. Nutrition Journal, 10(1):26, 2011.

[19] René W. M. M. Jansen and Lewis A. Lipsitz. Postprandial hypotension: epidemiology, pathophysiology, and clinical management. Annals of Internal Medicine, 122(4):286–295, 1995. PMID: 7825766.

[20] K. L. Jones, C. MacIntosh, Y. C. Su, F. Wells, I. M. Chapman, A. Tonkin, and M. Horowitz. Guar gum reduces postprandial hypotension in older people. Journal of the American Geriatrics Society, 49(2):162–167, 2001.

[21] Claude Julien. The enigma of mayer waves: facts and models. Cardiovascular research, 70(1):12–21, 2006.

[22] Mark T. Kearney, Alan J. Cowley, and Ian A. Macdonald. The cardiovascular responses to feeding in man. Experimental Physiology, 80:683–700, 1995.

[23] H. Kelbæk, O. Munck, N. J. Christensen, and J. Godtfredsen. Central haemodynamic changes after a meal. British Heart Journal, 61(6):506–509, 1989.

[24] Henning Kelbæk, Thomas GjØrup, Niels Juel Christensen, Ole Munck, and John Godtfredsen. Central hemodynamic changes after ingestion of a meal in patients with coronary artery disease. Archives of internal medicine, 149(2):363–365, 1989.

[25] Tomasz Krauze, Małgorzata Liberkowska, Karolina Adamska, Elżbieta Turska, Andrzej Wykrętowicz, and Przemysław Guzik. Acute hemodynamic effects of salted potato chips in healthy people. Polish Archives of Internal Medicine, 2019.

[26] N. Lipovetzky, H. Hod, A. Roth, Y. Kishon, S. Sclarovsky, and M. S. Green. Heavy meals as a trigger for a first event of the acute coronary syndrome: a case-crossover study. Israel Medical Association Journal, 6:728–731, 2004.

[27] Ching-Liang Lu, Xiaoping Zou, William C Orr, and JDZ Chen. Postprandial changes of sympathovagal balance measured by heart rate variability. Digestive diseases and sciences, 44(4):857–861, 1999.

[28] MD Gina L Luciano, MD Maura J Brennan, and MD Michael B Rothberg. Postprandial hypotension. The American journal of medicine, 123(3):281–e1, 2010.

[29] D. B. MacLeod, L. I. Cortinez, J. C. Keifer, D. Cameron, D. R. Wright, W. D. White, et al. The desaturation response time of finger pulse oximeters during mild hypothermia. Anaesthesia, 60(1):65–71, 2005.

[30] Gregory L. Moneta, David C. Taylor, W. Scott Helton, Michael W. Mulholland, and D. Eugene Strandness, Jr. Duplex ultrasound measurement of postprandial intestinal blood flow: effect of meal composition. Gastroenterology, 95(5):1294–1301, 1988.

[31] Magnus Nordin. Sympathetic discharges in the human supraorbital nerve and their relation to sudo- and vasomotor responses. The Journal of physiology, 423(1):241–255, 1990.

[32] D. O’Donovan, C. Feinle-Bisset, C. Chong, A. Cameron, A. Tonkin, J. Wishart, M. Horowitz, and K. L. Jones. Intraduodenal guar attenuates the fall in blood pressure induced by glucose in healthy older adults. Journals of Gerontology Series A: Biological Sciences and Medical Sciences, 60(7):940–946, 2005.

[33] Michael F. O’Rourke and Alfredo L. Pauca. Augmentation of the aortic and central arterial pressure waveform. Blood Pressure Monitoring, 9(4):179–185, 2004. PMID: 15311144.

[34] Stephen Payne. Cerebral autoregulation: control of blood flow in the brain, volume 15. Springer, 2016.

[35] Karen Reminy, Olivier Hue, and Sophie Antoine-Jonville. Effect of warm environment on the skin blood flow response to food intake. International Journal of Hyperthermia, 37(1):836–842, 2020.

[36] A. Russo, J. E. Stevens, T. Wilson, F. Wells, A. Tonkin, M. Horowitz, and K. L. Jones. Guar attenuates fall in postprandial blood pressure and slows gastric emptying of oral glucose in type 2 diabetes. Digestive Diseases and Sciences, 48:1221–1229, 2003.

[37] M. B. Sidery and I. A. Macdonald. The effect of meal size on the cardiovascular responses to food ingestion. British Journal of Nutrition, 71:835–848, 1994.

[38] M. B. Sidery, I. A. Macdonald, and P. E. Blackshaw. Superior mesenteric artery blood flow and gastric emptying in humans and the differential effects of high fat and high carbohydrate meals. Gut, 35:186–190, 1994.

[39] Michael B. Sidery, Ian A. Macdonald, Alan J. Cowley, and L. J. Fullwood. Cardiovascular responses to high-fat and high-carbohydrate meals in young subjects. American Journal of Physiology, 261(5 Pt 2):H1430–H1436, 1991.

[40] K. Takazawa, N. Tanaka, M. Fujita, O. Matsuoka, T. Saiki, M. Aikawa, et al. Assessment of vasoactive agents and vascular aging by the second derivative of photoplethysmogram waveform. Hypertension, 32:365–370, 1998.

[41] Jennifer L. Taylor, Timothy B. Curry, Luke J. Matzek, Michael J. Joyner, and Darren P. Casey. Acute effects of a mixed meal on arterial stiffness and central hemodynamics in healthy adults. American Journal of Hypertension, 27(3):331–337, 2014.

[42] N. Tentolouris, C. Tsigos, D. Perea, E. Koukou, D. Kyriaki, E. Kitsou, S. Daskas, Z. Daifotis, K. Makrilakis, S. A. Raptis, and N. Katsilambros. Differential effects of high-fat and high-carbohydrate isoenergetic meals on cardiac autonomic nervous system activity in lean and obese women. Metabolism, 52(11):1426–1432, 2003.

[43] Laurence G Trahair, Lora Vanis, Diana Gentilcore, Kylie Lange, Christopher K Rayner, Michael Horowitz, and Karen L Jones. Effects of variations in duodenal glucose load on blood pressure, heart rate, superior mesenteric artery blood flow and plasma noradrenaline in healthy young and older subjects. Clinical Science, 122(6):271–279, 2012.

[44] Marleen A. van Baak. Meal-induced activation of the sympathetic nervous system and its cardiovascular and thermogenic effects in man. Physiology & Behavior, 94(2):178–186, 2008.

[45] B. A. Waaler, M. Eriksen, and K. Toska. The effect of meal size on postprandial increase in cardiac output. Acta Physiologica Scandinavica, 142(1):33–39, 1991.

[46] I. B. Wilkinson, H. MacCallum, L. Flint, J. R. Cockcroft, D. E. Newby, and D. J. Webb. The influence of heart rate on augmentation index and central arterial pressure in humans. The Journal of Physiology, 525(Pt 1):263–270, 2000.

[47] Walter C Willett, Geoffrey R Howe, and Lawrence H Kushi. Adjustment for total energy intake in epidemiologic studies. The American journal of clinical nutrition, 65(4):1220S–1228S, 1997.

[48] World Health Organization. Obesity: preventing and managing the global epidemic. Report of a WHO consultation. WHO Technical Report Series 894, World Health Organization, Geneva, 2000.

[49] Rui Zhang and Oliver Amft. Monitoring chewing and eating in free-living using smart eyeglasses. IEEE journal of biomedical and health informatics, 22(1):23–32, 2017.

